# Stressful times in a climate crisis: how will aphids respond to more frequent drought?

**DOI:** 10.1101/2020.06.24.168112

**Authors:** Daniel Joseph Leybourne, Katharine F Preedy, Tracy A Valentine, Jorunn IB Bos, Alison J Karley

## Abstract

**Aim:** Aphids are abundant in natural and managed vegetation, supporting a diverse community of organisms and causing damage to agricultural crops. Using a meta-analysis approach, we aimed to advance understanding of how increased drought incidence will affect this ecologically and economically important insect group, and to characterise the underlying mechanisms.

**Location:** Global.

**Time period:** 1958–2020.

**Major taxa studied:** Aphids.

**Methods:** We used qualitative and quantitative synthesis techniques to determine whether drought stress has a negative, positive, or null effect on aphid fitness. We examined these effects in relation to 1) aphid biology, 2) the aphid-plant. species combination. We compiled two datasets: 1) a “global” dataset (*n* = 55 from 55 published studies) comprising one pooled effect size per study, and 2) an “expanded” dataset (*n* = 93) containing multiple datapoints per study, separated into different measures of aphid fitness but pooled across aphid-plant combinations. Where reported, we extracted data on the effect of drought on plant vigour, and plant tissue concentrations of nutrients and defensive compounds, to capture the potential causes of aphid responses.

**Results:** Across all studies (“global” dataset), drought stress had a negative effect on aphid fitness: Hedges’ g = −0.57; 95% confidence interval (CI_95_) = ±0.31. The “expanded” dataset indicated that, on average, drought stress reduced aphid fecundity (g = − 0.98; CI_95_ = ±0.50) and increased development time (g = 1.13; CI_95_ = ±1.02). Furthermore, drought stress had a negative impact on plant vigour (g = −7.06; CI_95_ = ±2.86) and increased plant concentrations of defensive chemicals (g = 3.14; CI_95_ = ±3.14).

**Main conclusions:** Aphid fitness is typically reduced under drought, associated with reduced plant vigour and increased chemical defence in drought-stressed plants. We propose a conceptual model to predict drought effects on aphid fitness in relation to plant vigour and defence.

## Introduction

The changing climate is anticipated to lead to decreased annual levels of precipitation in some regions, resulting in extended periods of drought (Blenkinsop & Fowler 2007; Santos, Belo-Pereira, Fraga & Pinto 2016). For plants in mesic habitats, prolonged drought can have severe consequences on plant physiology, often leading to reduced growth and photosynthetic capacity (Osakabe, Osakabe, Shinozaki & Trans 2014; Zeppel, Wilks & Lewis, 2014). Plant physiological responses to drought can directly influence the population dynamics, fitness, phenology, and biology of herbivorous insects (Stayley et al., 2006; Huberty & Denno 2004; Mody, Eichenberger & Dorn, 2009; Aslam, Johnson & Karley, 2013), with consequences that cascade through trophic networks (Johnson, Stayley, McLeod & Hartley, 2011; Rodríguez-Castañeda 2013). Meta-analysis provides a useful approach to predict the direction of drought effects on insect herbivores, reporting an overall response that accommodates between-study variation.

Previous meta-analyses have examined drought effects by comparing responses of herbivorous insect species with different feeding strategies (Koricheva & Larsson, 1998; Huberty & Denno 2004). To date, however, there has been no comprehensive assessment of drought effects on a specific herbivore group and the underpinning causes due to physiological changes in the host plant. Aphids are phloem-feeding insects of global ecological importance (Van Emden & Harrington 2017) as abundant components of insect communities in diverse ecosystems across the globe (Messelink, Sabelis & Janssed, 2012; Roubinet et al., 2018). There are over 4,400 known species of aphid (Blackman & Eastop 2000) and around 250 of these are major agricultural and horticultural pests, making them an economically important group of herbivorous insects. In many ecosystems, aphids sustain several higher trophic groups, including their primary consumers, such as parasitoid wasps, spiders, ladybirds, and carabid beetles (Staudacher, Jonsson & Traugott, 2016), the higher-level consumers of these aphid natural enemies, such as hyperparasitoids (Traugott et al., 2008; Lefort et al., 2017), small mammals and birds, and many entomological pathogens and parasites (Hagen & van den Bosch 1968). Examining how climate change, including drought, might influence aphid fitness is a major avenue of current research, specifically with regards to examining how this might affect the productivity and functioning of agricultural, horticultural, and natural vegetation systems across the globe (Romo & Tylianakis 2013; Teixeira, Valim, Oliveira & Campos, 2020)

Analysis of sap-feeding insects by Huberty & Denno (2004) suggested that drought has an overall negative effect on the fitness of sap-feeding insects. Experimental studies of aphids indicate that this negative effect of drought is observed across many aphid-plant systems (Pons & Tatchell 1995; Agele, Ofuya & James, 2006; Mody et al., 2009; Aslam et al., 2013; Grettenberger & Tooker 2016; Foote, Davis, Crowder, Bosque-Pérez & Eigenbrode, 2017), although this has not been assessed quantitatively. Further, there has been no comprehensive analysis of the causes of decreased aphid fitness under drought, although several studies suggest that it is mediated through reduced plant fitness (Hale, Bale, Pritchard, Masters & Bworn, 2003; Banfield-Zanin & Leather, 2015; Dai, Liu & Shi, 2015). Two meta-analyses conducted in recent decades provide context for constructing a hypothesis to explain variation in aphid fitness under water stress in relation to plant fitness. First, Huberty & Denno (2004) found little evidence for the plant stress hypothesis (i.e. enhanced insect performance on water-stressed host plants due to increased tissue nitrogen availability: White, 1969) amongst sap-feeding insects (phloem and mesophyll feeders). Second, Cornelissen, Fernandes & Vasconcellos-Neto (2008) examined insect fitness in relation to plant vigour and demonstrated that sap-feeding insects are more abundant and show increased fitness when feeding on more vigorously growing plants. These findings lead us to hypothesise that the effects of drought on aphid fitness are driven by decreased plant vigour, such as reduced plant growth rate or mass (Hatier et al., 2014), rather than stress-related changes in plant nutritional quality.

Although many studies have reported reduced aphid fitness when exposed to drought stressed hosts (Banfield-Zanin & Leather, 2015; Dai et al., 2015; Foote et al., 2017), studies have reported null (Mewis, Khan, Glawischnig, Schreiner & Ulrichs, 2012) and positive (Oswald & Brewer, 1997) effects. Multiple factors could explain these contrasting observations, including differences in aphid or plant biology. Indeed, in the study by Oswald & Brewer a positive effect of drought on aphid fitness was detected in the Russian wheat aphid, *Diuraphis noxia* (Mordvilko), and a negative effect was reported for the corn leaf aphid, *Rhopalosiphum maidis* (Fitch). Although both species are cereal-feeding aphids, *D. noxia* and *R. maidis* belong to two distinct aphid tribes, the Macrosiphini and the Aphidini, respectively (Kim & Lee, 2008; Choi, Shin, Jung, Clarke & Lee, 2018), raising the possibility that differences in aphid biology and/or life history could underlie the contrasting responses. Additionally, the specific aphid-plant combination could further influence the effects of drought on aphid fitness. For example, multiple aphid species exhibit contrasting responses to drought on a common host plant (Mewis et al.) and a single aphid species can display contrasting responses on related host plant species (Hale et al., 2003). These findings suggest that aphid responses to drought could be mediated by plant species-specific responses to drought (i.e. the availability of nutrients, the concentration of defensive compounds, resource allocation to new tissue). Understanding these mechanisms is necessary to predict the outcomes of plant-insect interactions under a changing climate.

Here, we take the novel approach of analysing data on aphid fitness and host plant physiology, using studies comparing drought with unstressed conditions, to examine the hypothesis that changes in aphid fitness are driven by the effects of drought on plant vigour. We predicted that reduced aphid fitness would be associated more strongly with decreased plant vigour (e.g. decreased mass, reduced growth) under drought than with changes in plant nutritional quality or defensive chemistry. Initially, we carry out a literature synthesis and take a “vote-counting” approach to qualitatively determine whether drought has an overall negative, positive, or null effect on aphid fitness. Following this, we use meta-analysis techniques to quantify these effects. Next, we extract data reporting on plant physiological responses to drought, including measurements of plant vigour, and tissue concentrations of plant nutrients and plant defensive compounds. This provides us with data that can be used, for the first time, to quantify drought effects on plant physiology in parallel with aphid fitness responses. A secondary aim of the meta-analysis was to explore patterns in aphid responses to drought in relation to 1) aphid tribe or aphid host range, and 2) the aphid-plant system (i.e. species combinations) to identify any common features of aphid biology that explain variation in aphid fitness responses to drought. The mechanistic understanding provided by our study allows the effects of drought on herbivore success to be anticipated for phloem-feeding insects across different habitats under future climatic conditions.

### Literature search and meta-analysis

#### Criteria for inclusion in analysis

The search terms “Drought” AND “Aphid” were used to conduct a literature search of both the Web of Science and Scopus databases (with a publication cut-off date of September 2020), to maximise the number of studies included (the overlap between them is only 40-50%: Nakagawa, Noble, Senior & Lagisz, 2017). 190 papers were identified from the Web of Science database and 197 from the Scopus database. After removing duplicates, 247 published papers were extracted. A previous meta-analysis which examined insect responses to drought (Huberty & Denno, 2004) was screened and an additional 16 studies were identified. This produced a pool of 263 studies published between 1958 and 2020. See Fig. 1 for the PRISMA flow diagram, constructed following Moher, Liberati, Tetzlaff & Altman (2009). To be considered for inclusion in the analysis, papers had to satisfy the following criteria: 1) to be primary literature presenting data on the responses of at least one aphid species to drought relative to an unstressed control treatment; 2) report aphid responses as the effect of drought on a measure of aphid fitness; 3) present the responses so that an estimation of the treatment differences could be determined alongside an estimate of the variation. A total of 55 studies satisfied these criteria. A further 26 studies reported data for aphid fitness but did not display the data; these studies were excluded from the meta-analysis but included in qualitative “vote-counting” assessment. The full range of studies are detailed in Appendix 1 and Appendix 2.

**Fig. 1:**
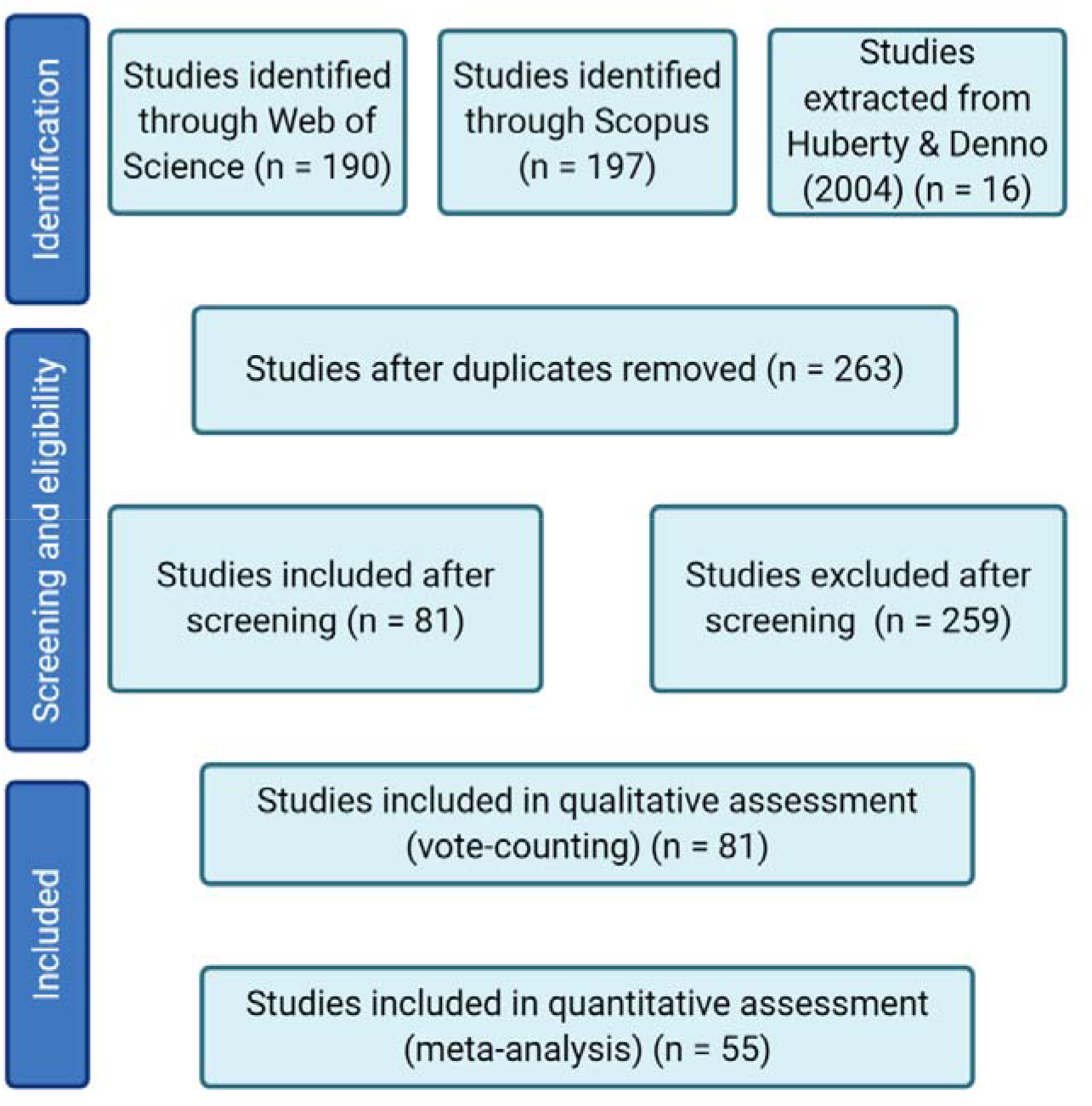
PRISMA diagram

Heterogeneity is routinely expected and accepted in meta-analyses (Higgins, 2008). Acceptance of heterogeneous data is dependent on whether the inclusion criteria are sound and the underlying data correct (Higgins). Extracted data were assessed for heterogeneity by measuring Cochran’s Q.

#### Data extraction and pooling: aphid responses

Aphid fitness data were extracted from drought and control (unstressed) treatments. Where reported, the mean value and an indication of the variation around the mean were extracted using WebPlotDigitizer v.4.2 (Rohatgi, 2019. Weblink: https://automeris.io/WebPlotDigitizer). Where median and interquartile ranges were reported, means and standard deviation were estimated following Luo, Wan, Liu & Tong (2018) and Wan, Wang, Liu & Tong (2014). Data were extracted for the following aphid fitness parameters: fecundity (daily, lifetime, and mean fecundity and life-history parameters related to reproduction, such as the intrinsic rate of increase), population size or aphid density/abundance, aphid development (time until adulthood and time until first reproduction), aphid biomass, or aphid lifespan. The effect size (Hedges’ g; Cooper, Hedges, Valenting, 2019) was calculated from aphid responses under drought relative to aphid responses under control conditions. Hedges’ g was selected as this parameter performs better across studies with a lower sample size, when compared with Cohen’s D or Glass’s Delta. Where multiple drought treatments were imposed, data were extracted from the control and the most severe drought treatment.

Where multiple formats of a fitness parameter were reported (e.g. fecundity reported in terms of mean fecundity and lifetime fecundity) data were pooled across different measures of the core parameter to provide one response per parameter assessed. Data were further pooled across any other experimental treatments imposed in the study (for example, in Xing et al. (2003) drought and control conditions were pooled across three CO_2_ treatments), and within aphid species to provide one data point per aphid species per fitness parameter. Data were collated separately for each host plant species. This pooling method produced 93 unique data points over the 55 studies.

Two datasets were compiled based on these data. In the “global” dataset (see Supplementary Fig. 1A for the associated funnel plot), data were pooled within each study across fitness parameters, plant hosts, and aphid species to produce 55 data points, i.e. one pooled effect size per study, reporting the overall aphid responses to drought. Within the “global” dataset, the calculated hedges’ g values for developmental fitness parameters were converted from positive to negative values to align with the net direction of other fitness parameters (this was required because a positive hedges’ g value for development represents a fitness decrease, compared with the other parameters for which a fitness decrease would result in a negative hedges’ g value). In the “expanded” dataset (see Supplementary Fig. 1B for the associated funnel plot), effect sizes were calculated separately for each fitness parameter and aphid-host plant combination in the study. To ensure that the direct comparison of different aphid fitness parameters was justified for inclusion in the analysis, the calculated hedges’ g values for each data point were plotted (Supplementary Fig. 2) to confirm that data were evenly distributed and not clustered into categories. Furthermore, to avoid potential pseudo-replication resulting from extracting multiple fitness parameters per study, the study number was included as a random term in all analyses of the “expanded” dataset.

#### Grouping of extracted data: aphid responses

Extracted data contained information on 23 aphid species (Table 1). To disentangle any potential taxonomic differences, data were categorised into three groups: a “Tribe” grouping based on the taxonomic tribe of the aphid species; an “Aphid Host Range” category based on whether the aphid species is a specialist or generalist feeder; and a “Plant-Aphid” group based on the plant-aphid system examined. Categorisations for Tribe and Aphid Host Range are detailed in Table 1. The final Tribe grouping used in the “expanded” dataset comprised Macrosiphini (*n* = 60), Aphidini (*n* = 23), and others (*n* = 10), while the aphid host range grouping comprised Specialist (*n* = 62) and Generalist (*n* = 31) categories.

**Table 1:** information on the aphid species included in the meta-analysis, the agricultural and ecological importance of each species, and the number of data points present in the “Expanded” dataset.

| Aphid species | Common name | Aphid tribe /<br>Specialism<br>(generalist or<br>specialist) | Agricultural and ecological<br>importance | Number of data<br>points: “Expanded”<br>dataset and study<br>from which data were<br>extracted □ |
| --- | --- | --- | --- | --- |
| *Adelges abietis*<br>Linnaeus | Eastern spruce gall<br>aphid | Adelgini (Other) /<br>Specialist | Common pest of Norway and Sitka spruce.<br>Stem-mother forms pineapple-shaped gall. | 2 datapoints from studies 10<br>and 11 |
| *Aphis glycines*<br>Matsumura | Soybean aphid | Aphidini / Specialist | Significant pest of soybean across North<br>America and Asia | 1 datapoint from study 31 |
| *A. Pomi* DeGeer | Apple aphid | Aphidini / Specialist | Significant pest of apple trees, specifically<br>on nursery stock. Colonies can be<br>attended by ants | 1 datapoint from study 29 |
| *Acyrtosiphon pisum*<br>(Harris) | Pea aphid | Macrosiphini /<br>Generalist | Widespread in temperate regions. Serious<br>pest of the Fabaceae. Vector of over 30<br>plant viruses | 4 datapoints from studies 7,<br>20, 26, 43 |
| *Brevicoryne brassicae*<br>(Linnaeus) | Cabbage aphid | Macrosiphini /<br>Generalist | Wax-covered aphid. Widely distributed<br>throughout Europe, major pest of<br>Brassicaceae | 8 datapoints from studies<br>23, 37, 47, 48, 49, |
| *Cinara costata*<br>(Zetterstedt) | Mealy spruce aphid | Eulachnini (Other) /<br>Specialist | Wax-covered aphid. Can cause damage to<br>spruce trees. Unlike other *Cinara spp.* *C.*<br>*costata* is not readily attended by ants. | 1 datapoint from study 24 |
| *C. pinea* (Mordvilko) | Large pine aphid | Eulachnini (Other) /<br>Specialist | Forestry pest of Scotts pine and other<br>*Pinus spp.* Often found on young pine<br>trees. | 1 datapoint from 32 |
| *Chaitophorus*<br>*leucomelas* Koch | Black polar leaf aphid | Chaitophorini (Other) /<br>Specialist | A widely distributed pest of poplar.<br>Colonies can reside in the galls formed by<br>other insects and can occasionally be<br>attended by ants. | 1 datapoint from study 39 |
| *Diuraphis noxia*<br>(Mordvilko) | Russian wheat aphid | Macrosiphini /<br>Specialist | A significant cereal pest in South Africa<br>and North America. Vector of barley yellow<br>dwarf virus. | 3 datapoints from studies 2,<br>16, 34 |
| *Dysaphis plantaginea*<br>(Passerini) | Rosy apple aphid | Macrosiphini /<br>Specialist | Widely distributed across temperate<br>regions and a significant pest of apples.<br>Colonies develop in galls and are attended<br>by ants. | 1 datapoint from study 41 |
| *Elatobium abietinum*<br>(Walker) | Green spruce aphid | Macrosiphini /<br>Specialist | Distributed across Europe and North<br>America. Can cause economic and<br>environmental damage by causing needle<br>defoliation on Spruce trees. | 6 datapoints from studies 2,<br>4, 5, 6 |
| *Lipaphis erysimi*<br>(Kaltenbach) | Wild crucifer aphid | Macrosiphini /<br>Specialist | Can feed on various Brassicaceae crops. | 1 datapoint form study 23 |
| *Macrosiphum euphorbiae* (Thomas) | Potato aphid | Macrosiphini / Generalist | A significant pest species with worldwide distribution. Can cause significant economic damage through the transmission of over 20 plant viruses. | 7 datapoints from studies 8, 33, 40 |
| *Metopolophium festucae* subsp. *Cerealium* (Theobald) | Fescue aphid | Macrosiphini / Specialist | A subspecies of *M. festucae* which can be a significant pest of cereals and grasses. | 2 datapoints from study 18 |
| *Myzus persicae* (Sulzer) | Peach-potato aphid | Macrosiphini / Generalist | Significant pest with worldwide distribution. Broad host range. Can cause significant economic damage through the transmission of over 40 plant viruses. | 12 datapoints from studies 27, 35, 38, 44, 47, 48, 49, 50 |
| *Pemphigus betae* Doane | Sugarbeet root aphid | Hormaphidini (Other) / Specialist | Root-feeding aphid, a main economic pest of sugar beet. | 2 datapoints from study 30 |
| *Phloeomyzus passerinii* (Signoret) | Poplar woolly aphid | Phloeomyzinae (Other) / Specialist | Distributed in temperate regions. Significant economic pest of poplar. | 1 datapoint from study 14 |
| *Rhopalosiphum maidis* (Fitch) | Green corn aphid | Aphidini / Specialist | Widely distributed worldwide. A significant pest of cereals. Commonly attended by ants. Vector of several plant viruses. | 2 datapoints from studies 34, 42 |
| *R. padi* (Linnaeus) | Bird cherry-oat aphid | Aphidini / Specialist | Widely distributed worldwide, a significant pest of cereals and a vector of numerous plant viruses. Populations on the primary host can be attended by ants. | 18 datapoints from studies 3, 9, 15, 18, 19, 21, 28 53, 55 |
| *Sitobion avenae* Fabricius | Wheat aphid | Macrosiphini / Specialist | Widespread pest of cereals with worldwide distribution. A vector of several economically important plant viruses. | 16 datapoints from 1, 13, 17, 25, 36, 45, 46, 51, 54 |
| *Sipha maydis* Passerini | Bristly black grass aphid | Siphini (Other) / Specialist | Widely distributed pest that feeds on numerous Poaceae species, including cultivated crops and perennial grasses. Can be attended by ants. | 1 datapoint from 28 |
| *Schizaphis graminum* (Rondani) | Spring green aphid | Aphidini / Specialist | Widely distributed pest of cereals. Can vector a range of plant viruses. | 1 datapoint from study 12 |
| *Therioaphis trifolii* (f. *maculate*) (Buckton) | Spotted alfalfa aphid | Panaphidina (Other) / Specialist | Widespread pest of Fabaceae. | 1 datapoint from study 7 |

☐ Full study references are contained in Appendix 2

Extracted data covered 24 host plant species. To facilitate comparison between plant-aphid combinations, data were grouped into the following plant host categories: Brassicas (comprising *Brassica oleracea* and *B. napus*, Bn: *n* = 6 and *15* from the “global” and “expanded” datasets, respectively), Cereals (*Arrhenatherum elatius, Dactylis glomerate, Holcus lanatus, Hordeum vulgare, Triticum aestivum: n = 20* and *38*), Forage species (*Chenopodium sp., Festuca arundinacea, Medicago sativa, Lolium perenne, L. multiforum, Poa autumnalis, Rumex sp.:: n = 7* and *10*), Legumes (*Pisum sativum, Glycine max: n = 2* for both datasets), Model *Arabidopsis* species (*Arabidopsis thaliana: n = 3* and *4*), *Solanum sp*. (*Solanum tuberosum, S. lycopersicum: n = 4* and 9), and Tree systems (*Malus domestica, Picea abies, P. sitchensis, P. Sylvestris, Populus sp., n = 13* and *15*).

#### Criteria for inclusion in analysis, data pooling, and data grouping: plant responses

Studies were screened for inclusion in an additional meta-analysis to determine the impact of drought on the host plant. To be considered for inclusion, studies had to satisfy the following criteria: 1) present data on the responses of at least one plant species to drought relative to a controlled condition; 2) report responses as the effect of drought on either a measure of vigour (including mass, height, and growth), tissue nitrogen (N) or amino acid concentration, or plant chemical defence (e.g. secondary metabolite or phytohormone concentration); 3) report an estimation of the differences alongside the variation. From the pool of 55 studies, 32 reported effects of drought on plant vigour (i.e. dry matter accumulation, plant growth, and leaf/tiller production), 12 reported a measure of tissue N or amino acid concentration, and ten reported tissue defensive compound concentrations. The effect size (Hedges’ g) was calculated as described above. Data were pooled at the study level into measures of vigour, N or amino acid concentration, and defensive compound concentrations, resulting in three plant sub-datasets: vigour, nutritional, and defensive.

#### Measuring publication bias

Funnel plots were created to test for publication bias and a rank correlation test was carried out to test for funnel plot asymmetry (Supplementary Fig. 1). Additionally, as most null results go unpublished, failsafe analysis using Orwin’s method (Orwin, 1983) was employed to estimate the number of studies reporting a null effect that would be required to reduce the observed average effect size to −0.1. This analysis estimated *n = 261* and *n = 2228* null studies required to reduce the observed average effect size to −0.1 in the aphid and plant datasets, respectively.

#### Analysis of extracted data

Statistical analysis was carried out using R version 4.0.3, with additional packages ggplot2 v.3.3.2 (Wickham, 2016), meta v.4.15-1, and metafor v.2.4-0. A total of 81 and 55 studies were included in the vote counting and meta-analysis, respectively (detailed in Appendix 1 and Appendix 2).

To account for variation between different methodologies used to implement drought, drought methodology was allocated into one of five categories: FC (studies where % reduction in field capacity was used); DI (studies where decreased volume of irrigation was used); GM (studies which used a gravimetric method to adjust irrigation); CC (studies which used a calibration curve to help advise water irrigation regimes); and RW (studies where irrigation was simply restricted or withheld from the drought treated plants). The effect of these drought treatments on aphid fitness was examined to confirm that different methodologies used to initiate drought did not vary in their effects (Supplementary Fig. 3).

#### Aphid response analysis: Vote counting procedure

Briefly, studies were screened for whether a significant effect of drought on aphid fitness was detected, and whether the direction of the effect was positive or negative. Studies which reported non-significant results were categorised as null response. Data were deemed as significant based on the statistical reporting in each study (significance was determined by p = <0.05). For studies measuring the responses of several aphid species (such as Foote et al., 2017), the results of the statistical analysis at the drought treatment level, not the species x drought interaction level, were used to determine whether the observation was significant amongst all aphid species.

#### Aphid response analysis: Meta-analysis procedure

Two meta-analyses were carried out using the two datasets described above: 1) an analysis using the “global” dataset pooled across aphid species and host plant giving one pooled effect size per study (55 data points); 2) an analysis using the “expanded” dataset which was pooled at the aphid species and host plant levels and separated by aphid fitness parameter, giving multiple pooled effect sizes per study (93 data points).

For the first meta-analysis (“global” dataset), data were analysed using a linear mixed effects model fitted with restricted maximum likelihood distribution. The difference between the plant-aphid systems was examined in a subsequent model by incorporating this factor a fixed term. For the second meta-analysis (“expanded” dataset), ‘study’ was incorporated as a random term to account for multiple data points in some studies and to avoid potential pseudo-replication. Data were analysed in a similar way to the method above: briefly, a random linear mixed effects model fitted with restricted maximum likelihood distribution was used to examine differences between aphid tribes, aphid host range, and the plant-aphid system in three models. In a further model, the responses of the different aphid fitness parameters were tested.

#### Plant response analysis: Meta-analysis procedure

32, 12, and ten studies also reported the effect of drought on plant vigour, plant tissue nutrient concentration, and tissue defensive compound concentrations, respectively. Data were analysed using a random linear mixed effects model fitted with restricted maximum likelihood distribution. Data were analysed in an individual model for each plant response.

## Results

### Aphid fitness is reduced under drought

The vote counting procedure (Fig. 2) indicated that aphid fitness is reduced when exposed to drought stressed plants. Analysis of the pooled data indicated that aphid fitness is generally reduced under drought (Q_M_ = 26.71; p = <0.001; Supplementary Fig. 3), average Hedges’ g = −0.57; 95% confidence interval (CI_95_) = ±0.31. Subsequent comparisons were carried out after separating the data into the plant-aphid system. Pooled data were significantly heterogeneous, indicated by Cochran’s Q (Q_E_ = 404.34; df = 54; p = <0.001). Further comparison indicated that aphid fitness was broadly reduced under drought across plant-aphid systems (Q_M_ = 25.81; p = <0.001; Fig. 3). However, due to low replication for some groups, a comprehensive comparative analysis was not possible. No trends over publication time were observed for pooled aphid responses (Supplementary Fig. 8).

**Fig. 2:**
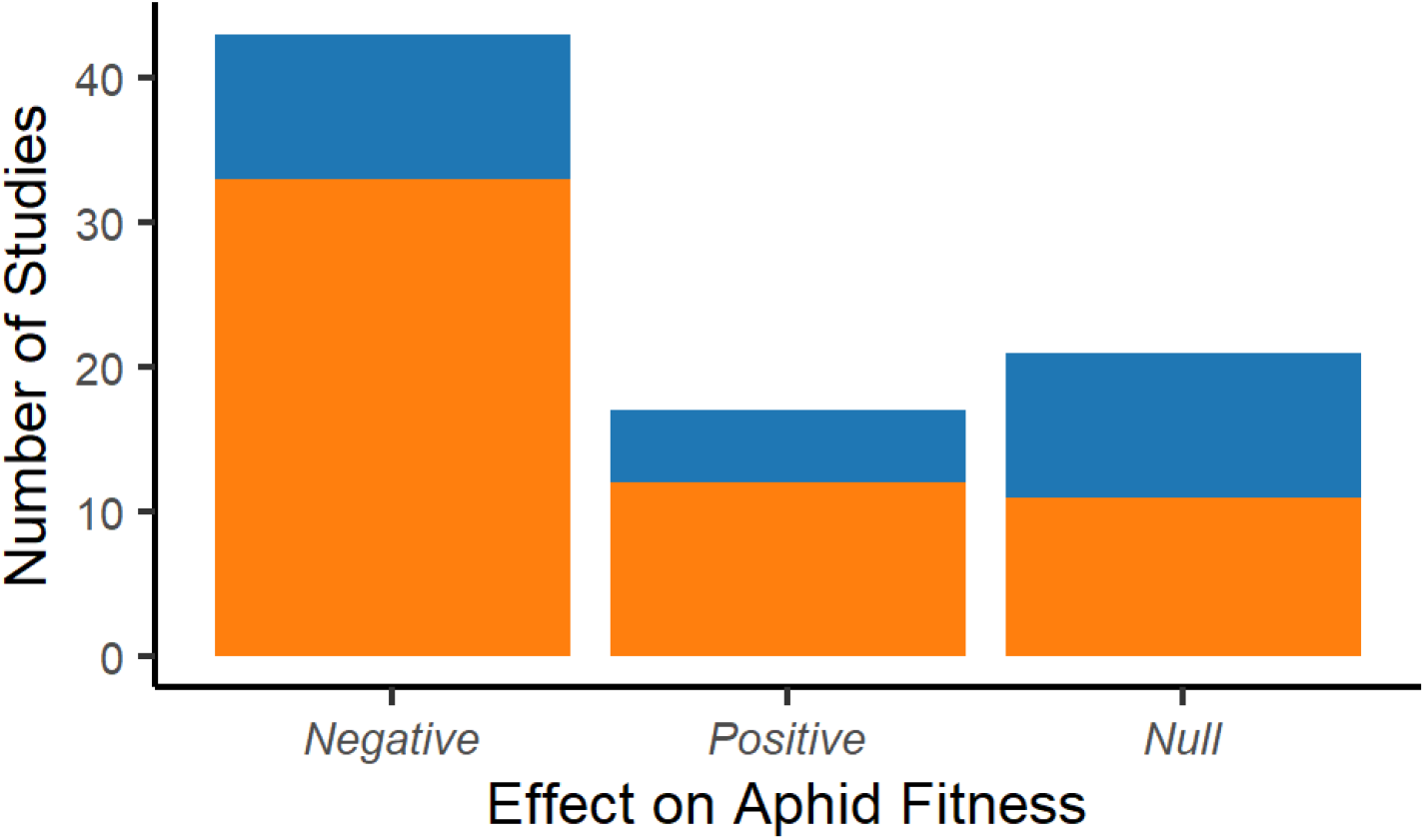
The number of studies reporting negative, positive, or null effects of drought stress on aphid fitness. Bars are coloured to indicate the proportion of studies in each category that were included in the full meta-analysis (Orange) vs those in the complete dataset (Blue).

**Fig. 3:**
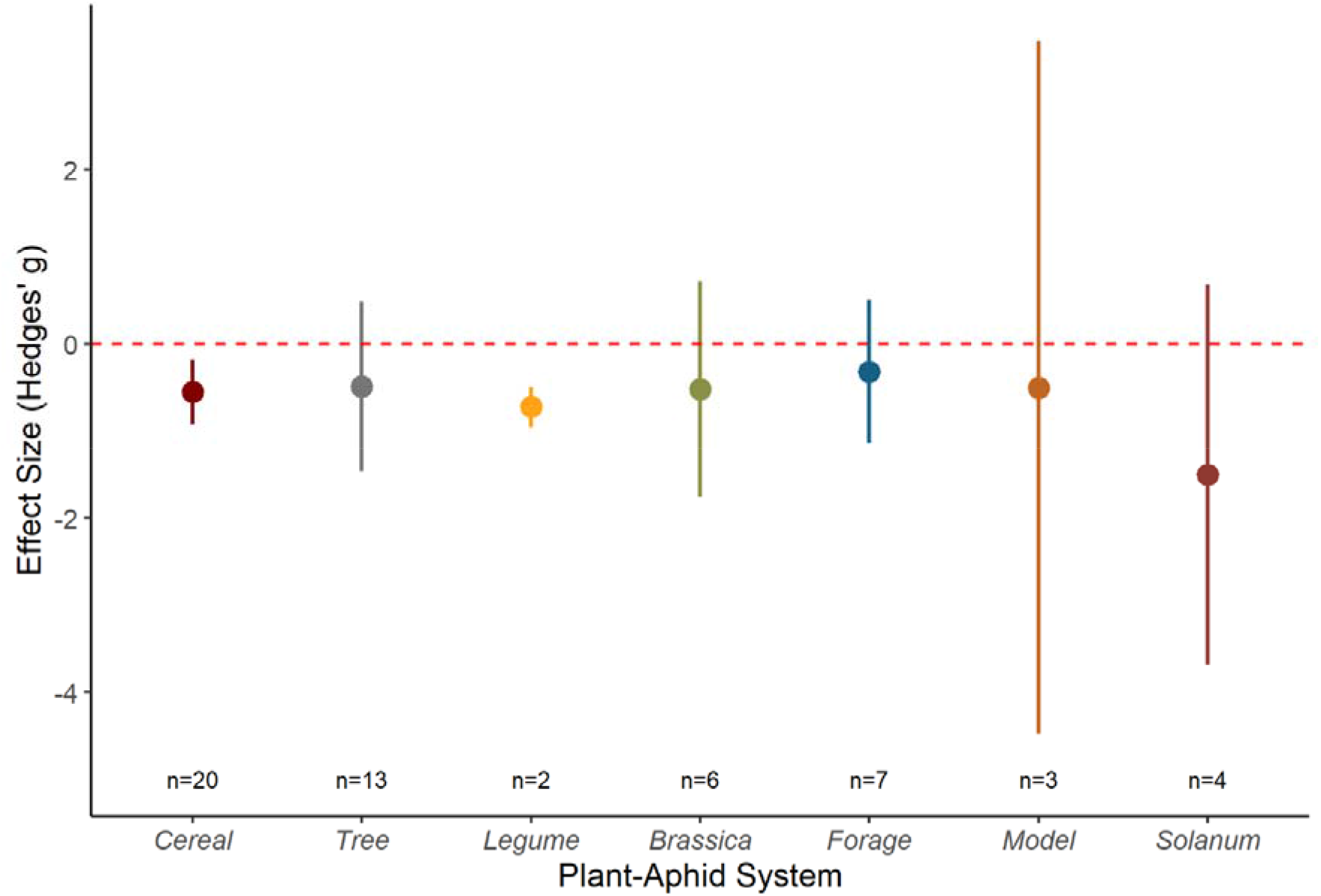
Effect of the plant-aphid system on overall aphid responses to drought stress conditions using the “global” dataset where response variables were pooled to produce one effect size per study. Graph displays the mean effect size (Hedges’ g) and the 95% confidence intervals for the different plant-aphid systems identified from the extracted data. Red dashed line represents zero effect size.

To analyse the effect of drought on aphid fitness in relation to aphid tribe and aphid host range, and to identify any differences between fitness parameters, a second dataset comprising 93 data observations over the 55 studies was compiled (the “expanded” dataset). Meta-analysis of this expanded dataset indicated that aphid fitness was reduced under drought (Q_M_ = 18.03; p = 0.003; Fig. 4 insert). Most aphid fitness parameters examined, especially parameters intrinsically associated with aphid abundance (fecundity and population size) and development, were negatively affected by drought (Q_M_ = 91.88; p = <0.001; Fig. 4). Drought decreased aphid fitness to a similar extent across the different aphid tribes (Q_M_ = 19.45; p = 0.002): Aphidini (g = −0.55; CI_95_ = ±0.38), Macrosiphini (g = −0.85; CI_95_ = ±0.37), and Other (g = −0.32; CI_95_ = ±1.18). Fitness decreases were similar for aphids with contrasting host plant ranges (Q_M_ = 14.13; p = 0.001): specialist (g = −0.67; CI_95_ = ±0.32), generalist (g = −0.90; CI_95_ = ±0.54).

**Fig. 4:**
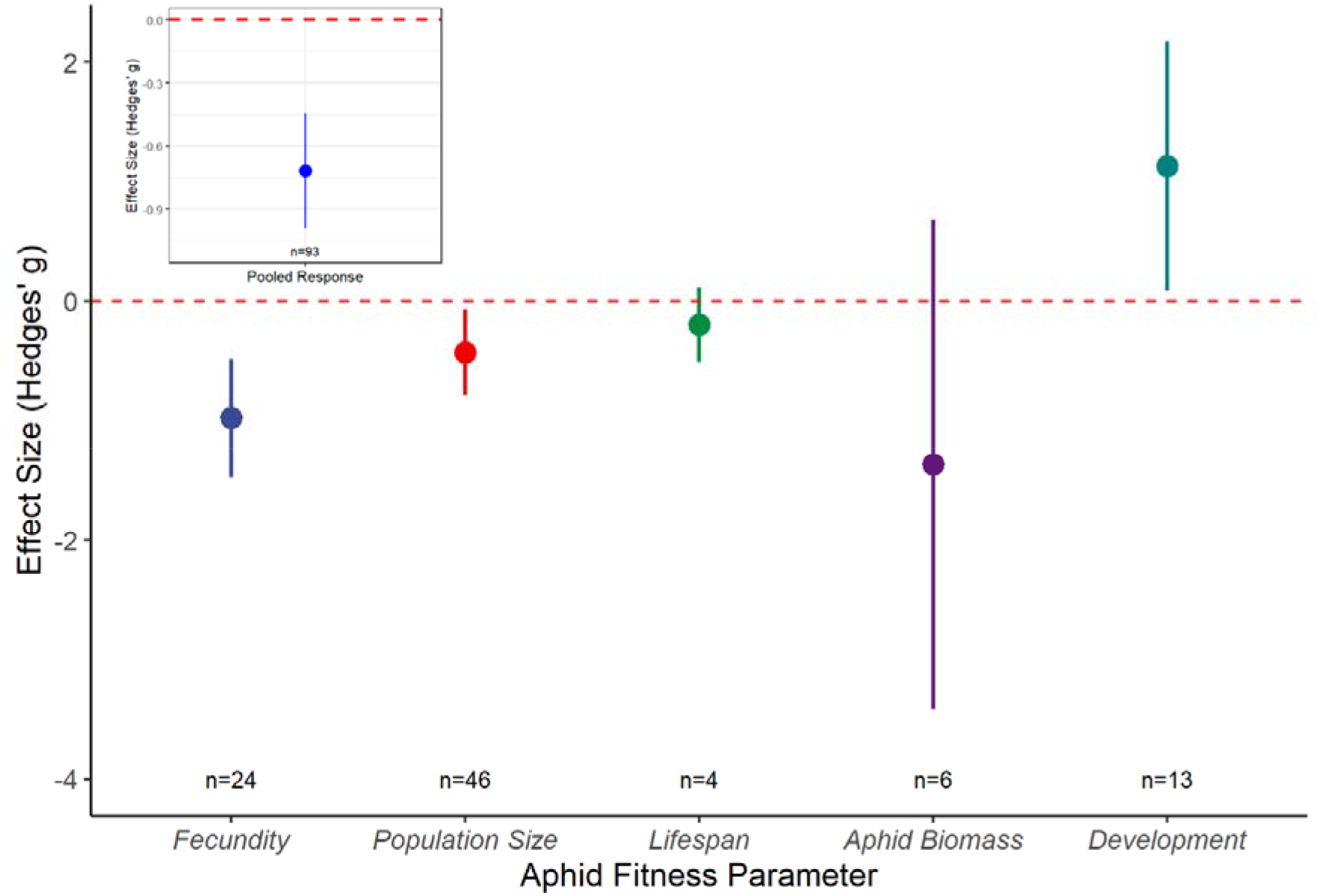
Responses of aphids to drought stress using the “expanded” dataset which included an effect size per response variable and for each aphid species measured in each study. Main graph displays the mean effect size (Hedges’ g) and the 95% confidence intervals for the different plant-aphid systems identified from the extracted data. Insert displays the mean effect size (Hedges’ g) and the 95% confidence intervals for the combined data. Red dashed line represents zero effect size.

### Drought stressed plants show reduced vigour and increased tissue concentrations of defensive compounds

Meta-analysis of the pooled plant responses to drought (Supplementary Fig. 5 – 7) indicated that exposure to drought has an overall negative effect on plant vigour (Q_M_ = 35.97; p = <0.001; Fig. 5) and, on average, results in more chemically-defended plant tissues (Q_M_ = 46.37; p = <0.001; Fig. 5). However, tissue N and amino acid concentrations did not increase consistently (Q_M_ = 4.92; p = 0.178; Fig. 5). The consistency of the relation between drought effects on aphid fitness, plant vigour and plant chemical defence, but not plant nutritional quality, is shown in Supplementary Fig. 9. No trends over publication time were observed for any plant responses (Supplementary Fig. 8).

**Fig. 5:**
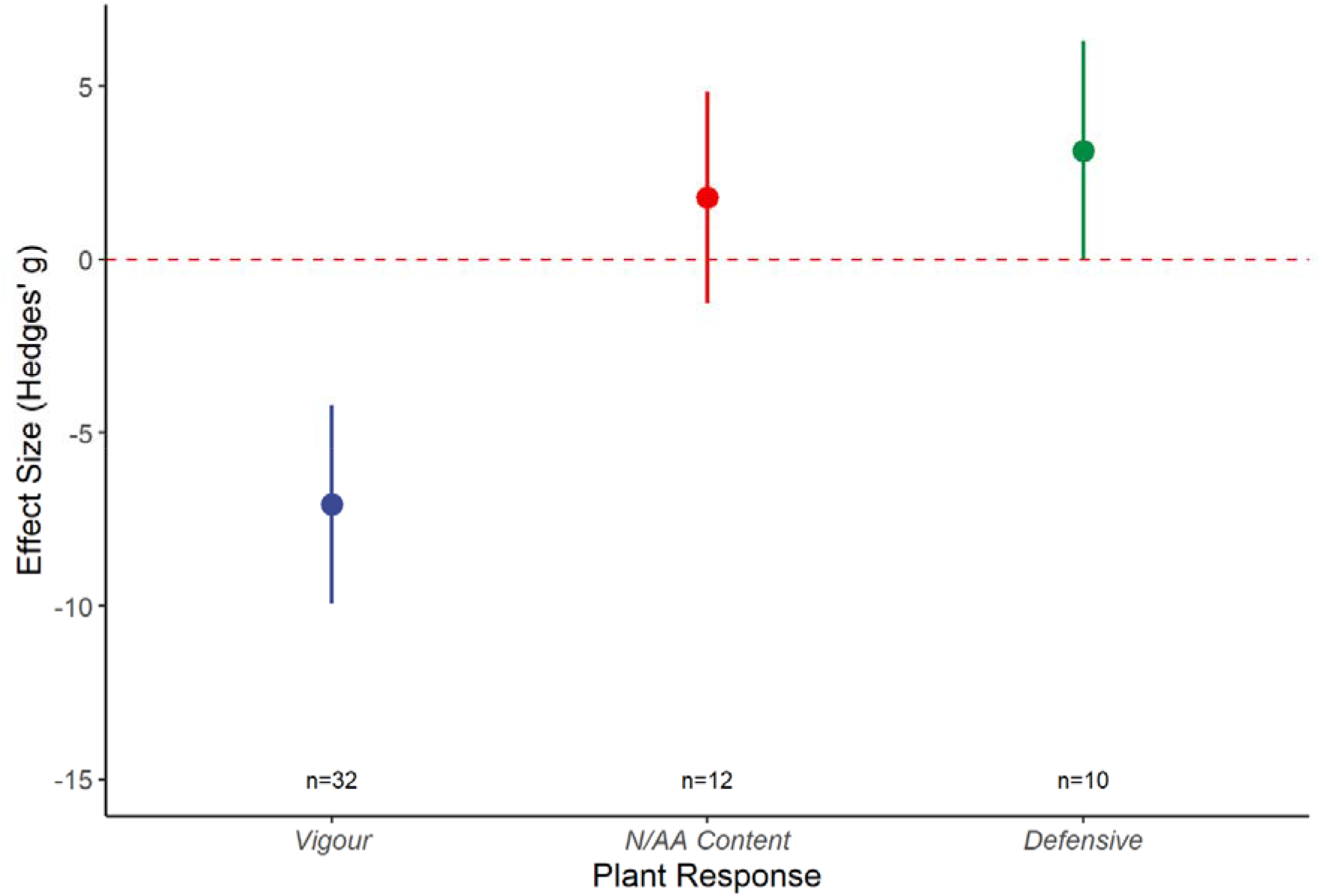
The effect of drought stress on plant vigour using the “plant” dataset where measures of plant vigour (vigour), N or amino acid (AA) concentration (N/AA Content), or tissue concentrations of defensive compounds (Defensive) were reported. Graph displays the mean effect size (Hedges’ g) and the 95% confidence intervals for the different plant-aphid systems identified from the extracted data. Red dashed line represents zero effect size.

## Discussion

This study provides the first comprehensive assessment of aphid and host plant responses to drought stress, analysed in terms of aphid fitness and plant vigour, nutritional quality, and chemical defence. The meta-analysis supported our prediction that drought reduces aphid fitness, and this effect was linked most consistently with reduced plant vigour rather than altered tissue nutrient concentrations, although increased chemical defence of plant tissues might also play a role. Our study includes data extracted from 81 (qualitative assessment) and 55 studies (quantitative assessment), including 42 papers for quantitative assessment published between 2003 – 2020 that were not included in an earlier meta-analysis across different insect feeding guilds (Huberty & Denno, 2004). Our findings, therefore, represent a significant advance in knowledge on the effects of drought on sap-feeding insects with a novel focus on drought effects on aphids as an ecologically important insect group.

### Plant vigour could explain drought effects on aphid fitness

The primary finding that aphid fitness is reduced when feeding on drought stressed hosts confirms the findings of Huberty & Denno (2004) for sap-feeding insects. Our study goes further, however, by providing evidence for the underlying mechanisms. We show that decreased aphid fitness is most frequently associated with the negative effects of drought on plant growth (vigour), with some evidence that plant tissue concentrations of defensive chemicals or signalling compounds play an important role. It should be noted that, out of the studies included in our meta-analysis, over half measured plant vigour (32 studies) while only around one-fifth measured tissue concentrations of defensive compounds (*n* = 10). Most of these studies showed a strong association between decreased aphid performance, reduced plant vigour and increased tissue concentrations of plant defensive compounds, but no clear association with tissue nutrient concentrations. This finding is consistent with our prediction and is supported by a large body of literature reporting elevated chemical defence and decreased plant growth and vigour under drought (Templer et al., 2017; Beetge & Krüger 2019; Xie et al., 2020). As less vigorous plants have a higher concentration of biochemical defences, drought stress could lead to increased aphid exposure to plant defensive compounds, leading to reduced fitness due to feeding deterrence and decreased phloem ingestion.

A secondary aim of this meta-analysis was to examine the consistency of drought effects across aphid groups and aphid-plant systems to identify factors which could explain differential effects (Oswald & Brewer, 1997; Hale et al., 2003). All aphid tribes assessed showed decreased aphid fitness in response to drought and a similar response was observed when aphids were categorised as specialists or generalists. However, there was an overrepresentation of the Aphidini and Macrosiphini tribes (45 out of the 55 studies) because many empirical studies focus on agriculturally or ecologically important aphid species, which are widely represented in these two tribes (Kim & Lee 2008; Choi et al., 2018). This limits the extent to which differences between tribes can be detected and further experimental work would be needed to confirm consistent drought responses across aphid groups. For example, it could be hypothesised that aphid species which actively remobilise plant nutrients (e.g. *D. noxia;* see Sandström, Telang & Moran, 2000) are less affected by drought than species that are unable to maintain a sufficient supply of plant nutrients. Similarly, species that can sequester plant defensive chemicals (e.g. *Brevicoryne brassicae* (Linnaeus); see Kazana et al. 2007) might better tolerate increased concentrations of toxic plant chemicals in response to drought.

Similarly, aphid fitness was generally reduced on drought stressed plants across all plant groups assessed, but there was an overrepresentation of two plant groups (Cereal and Tree groups), which prevents firm conclusions. Nonetheless, these findings confirm that the effect of drought on herbivorous insects is primarily mediated by general changes in plant physiology, as indicated by Cornelissen et al. (2008).

### The role of host plant defence in aphid responses to drought

Although based on a relatively small number of studies, our analysis indicated that plant chemical defence is elevated under drought, and this correlates with reduced aphid fitness. Several studies highlight the effects of anti-herbivore plant resistance strategies on aphid fitness under benign conditions (Guerrieri & Digilio, 2008; Greenslade et al., 2016), but few studies have examined whether host plant resistance against aphids is modified under climate stress. From the 55 studies assessed here, only five included observations of aphid responses on both susceptible and (partially)-resistant plant types (Oswald & Brewer, 1997; Björkman 2000; Dardeau et al., 2015; Verdugo, Sauge, Lacroze, Francis & Ramirez, 2015; Guo et al., 2016) with one further study examining aphid responses on resistant plants only (Ramirez & Verdugo, 2009). Additionally, three studies compared aphid responses on drought tolerant and drought susceptible host plants (De Farias, Hopper, Leclant, 1995; Rousselin et al., 2018; Quandahor, Lin, Gou, Coulter & Liu, 2019) and only one study examined the interactive effects of aphid resistance and drought tolerance (Grettenberger & Tooker, 2016). Such a low level of representation means that the interactive effect between plant resistance and drought could not be investigated empirically using meta-analysis. Of these studies, four report similar findings: aphid fitness is reduced on both susceptible and resistant plant hosts (Oswald & Brewer; Björkman; Dardeau et al.; Guo et al.), with a smaller reduction in fitness on the resistant host plant than on the susceptible host plant (Oswald & Brewer; Dardeau et al.). These findings indicate that while aphid fitness is reduced on resistant plants under benign conditions, under drought it decreases to similarly low values on susceptible and resistant plants. This highlights a significant knowledge gap in our understanding of how plant resistance traits are affected by environmental stress, which is becoming increasingly important for successful pest management in changing climatic conditions.

To stimulate further research into this potentially important interaction between plant resistance, herbivore success, and climatic conditions, we propose a simplified conceptual model to predict how aphids will respond to drought in relation to altered host-plant resistance, termed the Plant Resistance Hypothesis (Fig. 6). This expands upon previous conceptual models, the plant stress hypothesis and plant vigour hypothesis, which do not consider the potential differences in plant susceptibility to herbivorous insect pests. This new model suggests that under benign conditions the basal level of aphid fitness will differ between the susceptible (high aphid fitness) and resistant (moderate-to-low aphid fitness) plant types, as will the level of plant defence: susceptible (low level of defence) *vs*. resistant (high level of defence). Drought is likely to cause a reduction in plant vigour and decreased plant palatability for the aphid, characterised by elevated concentrations of plant defensive compounds (Inbar, Doostdar & Mayer, 2001; Ozturk et al., 2002), leading to differential changes in the chemical defence of susceptible (from low to high concentration) and resistant (continually high concentration) plants. Differential levels of basal aphid fitness between the two plant types leads to differences in the extent to which aphid fitness is affected by drought depending on whether the host is a susceptible (from high fitness to low fitness) or resistant (from intermediate fitness to low fitness) plant type.

**Fig. 6:**
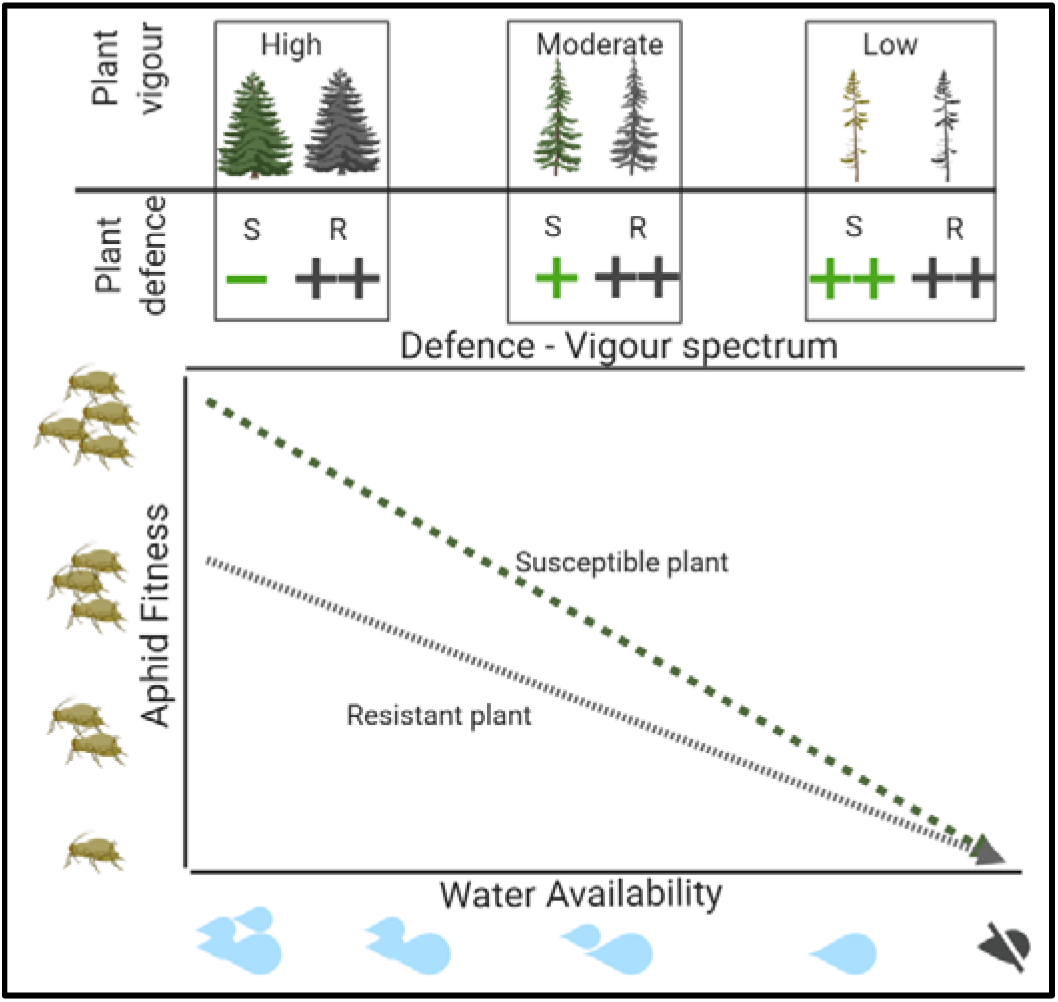
Conceptual representation of the Plant Resistance Hypothesis (PRH). As water availability decreases plant defence increases and plant vigour declines, leading to reduced aphid fitness. Basal levels of aphid fitness under conditions of ample water availability differ between the susceptible (S, green line) and the resistant (R, grey line) plant types. Under drought, aphid fitness is reduced on both plant types, however the extent of this reduction is greater for the susceptible plant (high to low fitness) than the resistant plant (intermediate to low fitness). This image made in BioRender © - biorender.com

### Drought-induced reduction in aphid fitness could destabilise aphid-trophic interactions

Aphids represent an important group of herbivorous insects from both an economic perspective, in relation to agricultural crop protection, and an ecological perspective, regarding the diverse community of higher trophic groups they support. The central finding of our meta-analysis is that exposure to drought-stressed hosts is detrimental to aphid fitness, although the extent of this effect is likely affected by host plant suitability as an aphid food source. Consequently, higher incidences of drought will have a detrimental effect on the terrestrial trophic networks that are supported by aphids: aphids are widespread in vegetation systems globally, are abundant consumers of primary production in many ecosystems, and provide a food source for many trophic groups (Gilbert 2005; Messelink et al., 2012; Roubinet et al., 2018). Our findings, largely based on aphid species in the Aphidini and Macrosiphini tribes, indicate that individual and population level measures of aphid fitness are affected negatively by drought, suggesting that drought will have cascading consequences for host plant consumption by aphids and the abundance of other trophic groups. If these findings translate to other aphid species and tribes, they imply that drought could reduce the availability of aphid hosts for parasites and pathogens (Nguyen, Michaud & Cloutier, 2007; Ahmed, Liu & Simon, 2012; Aslam, et al., 2013) and decrease food availability for aphid predators (Wade, Karley, Johnson & Hartley, 2017). Similarly, many aphids are tended by ants for their honeydew secretions (Stadler et al., 2003). These ants provide protective services to plants by deterring herbivory by other insect pests (Offenberg, Nielsen, MacIntosh, Havanon & Aksomkoae, 2004; Rosumek et al., 2009). While reduced aphid fitness might decrease plant consumption, it could also compromise the protective services delivered by ants: lower aphid abundance, or decreased honeydew quantity or quality, could decrease ant attendance (see Stadler, Kindlmann, Šmilauer & Fiedler, 2003), thereby exacerbating the detrimental effects of drought by increasing plant exposure to additional biotic stressors.

Ecological networks often exist in stable equilibria (McQuaid & Britton, 2015; Landi, Minoarivelo, Brännström, Hui & Dieckmann., 2018), with a change in the abundance of one species or functional group leading to perturbations in abundance and diversity across the network (McQuaid & Britton, 2015). A drought-induced reduction in aphid fitness might decrease abundance, mass, or quality of aphids available to support other trophic levels, with potential to destabilise population equilibria; a recent modelling study illustrated that the destabilising effects of drought on aphid-parasitoid interactions lead to altered insect population cycles (Preedy, Chaplain, Leybourne, Marion & Karley, 2020). A key finding of this meta-analysis was that plant resistance to aphids may influence the extent to which aphids are negatively affected by drought. The negative consequences of plant resistance for aphids generally include decreased fecundity (Greenslade et al., 2016; Leybourne et al., 2019) which could reduce aphid abundance for aphid natural enemies. This could have further consequences for aphid/insect distributions in regions that are experiencing more frequent drought events. Analysing the effects on aphid-natural enemy interactions of drought, and its interactions with other determinants of host suitability, is therefore an important avenue for future research to understand the impacts on the composition and function of ecological networks and species distributions (e.g. Rodríguez-Castaneda, 2013). This will improve our understanding of how drought and plant suitability characteristics might contribute to recent reports of increased rates of species loss and population declines (Saunders, Janes & O’Hanlon, 2020; Leather, 2018). Our conceptual model of the anticipated effects of drought on aphid fitness in relation to plant resistance, plant vigour, and chemical defence provides a basis for stimulating future research on insect-plant interactions under a changing climate.

## Data accessibility statement

Data will be uploaded to the Dryad data repository upon acceptance of the manuscript for publication.

## Appendices

## Appendix 1

References for the 25 studies which were included in the “vote-counting” analysis but excluded from the full meta-analysis

## Appendix 2

References for the 55 studies which were included in the full meta-analysis. □ indicates which studies contained more than one data point when split into the “expanded responses” dataset. * Indicates which studies reported on plant physiological, defensive, or nutritional responses for inclusion in the plant meta-analysis

