## Supplementary for "Stressful times in a climate crisis: how will aphids respond to more frequent drought?"

**Supplementary figures**

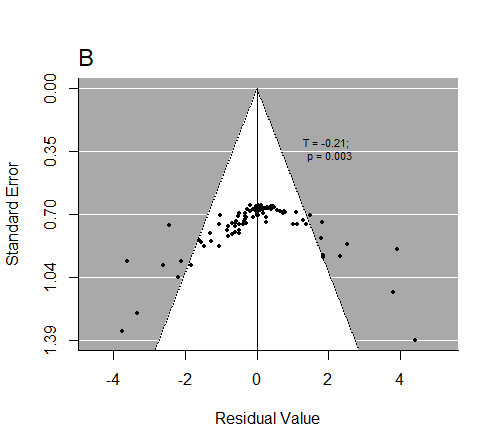

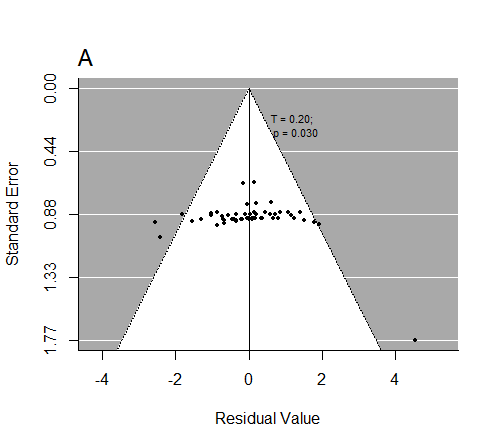

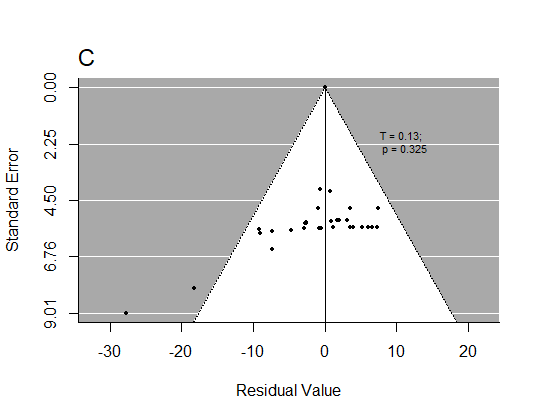

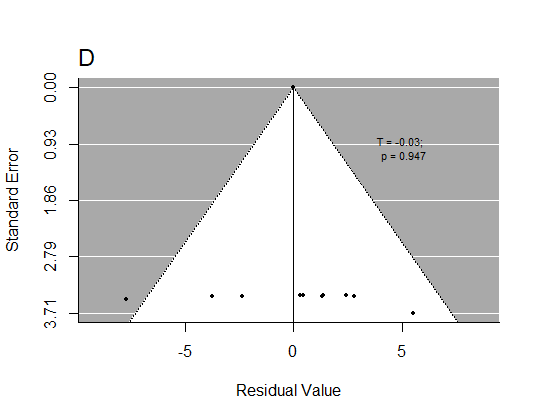

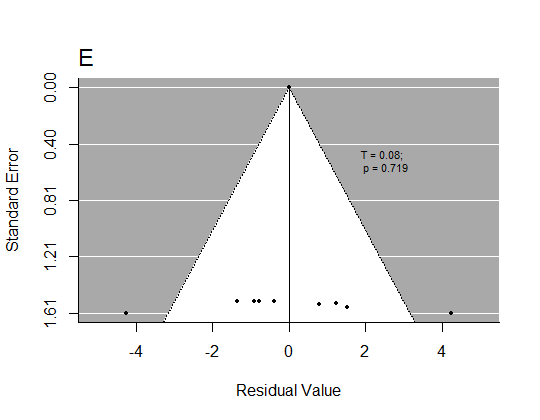

**Supplementary Fig. 1:** Funnel plots for the extracted studies. A) the “global” dataset, B) the “expanded” dataset, C) the dataset used in plant vigour analysis, D) the dataset used in plant nutrition analysis, E) the dataset used in plant defence analysis

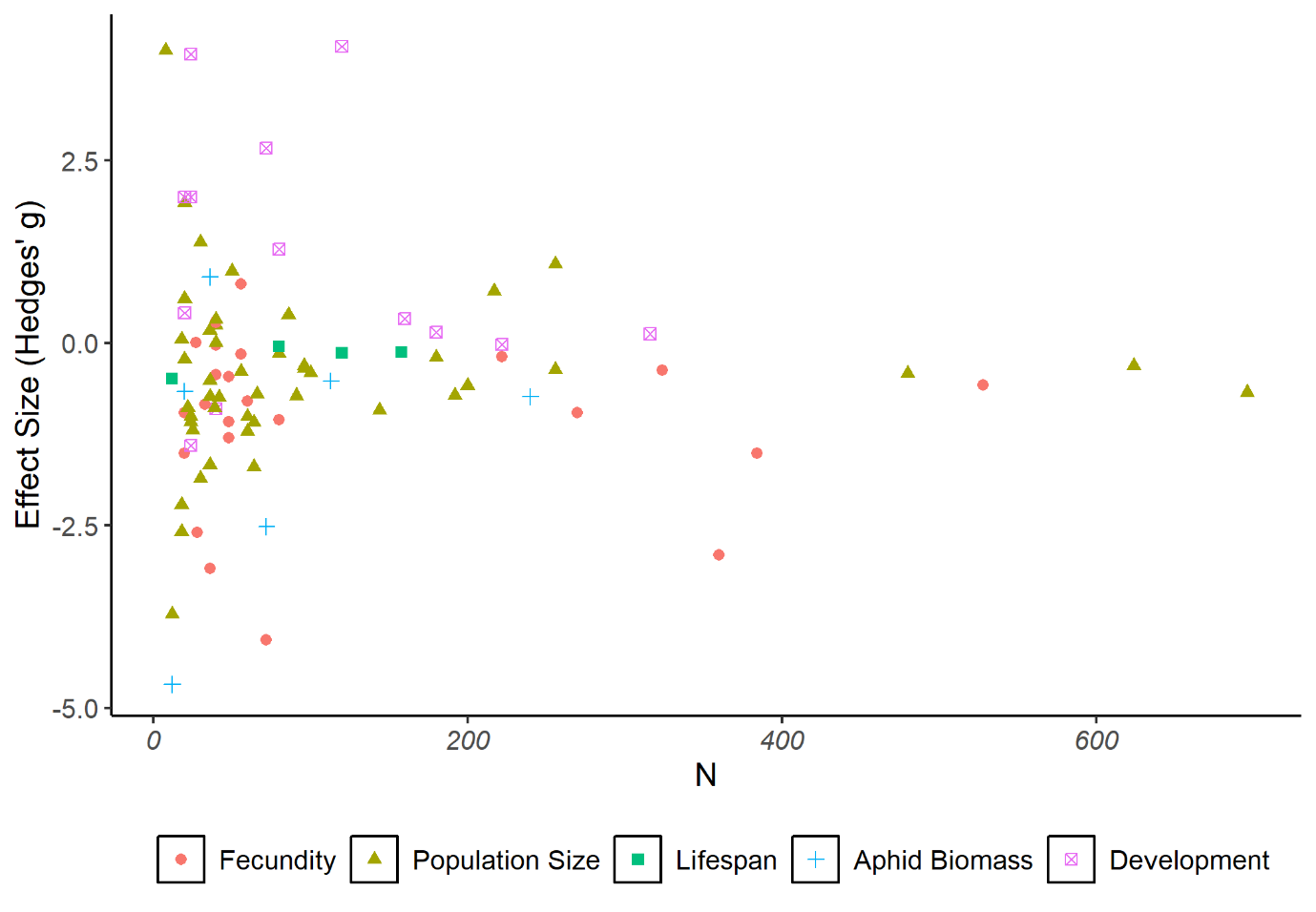

**Supplementary Fig. 2:** The observed effect size for each individual aphid fitness parameter in the “Expanded” dataset coded by the aphid fitness parameter it is categorised into. *N* represents the number of replicates (pooled across treatments to the drought treatment and aphid level) per observed point. Datapoint legend: filled circle – aphid biomass; filled triangle – development; filled square – fecundity; cross – lifespan; crossed open square – population size.

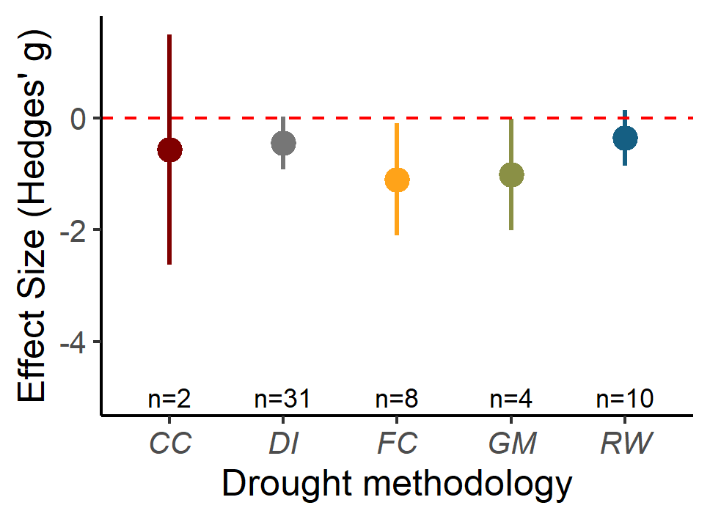

**Supplementary Fig. 3:** The effect of the methodology used to impose drought stress on pooled aphid fitness using the “global” dataset. Drought methodology coding: FC (studies where % reduction in field capacity was used); DI (studies where decreased volume of irrigation was used); GM (studies which used a gravimetric method to adjust irrigation); CC (studies which used a calibration curve to help advise water irrigation regimes); and RW (studies where irrigation was simply restricted or withheld from the drought treated plants). Graph displays the mean effect size (Hedges’ g) and the 95% confidence intervals for the different plant-aphid systems identified from the extracted data. Red dashed line represents zero effect size.

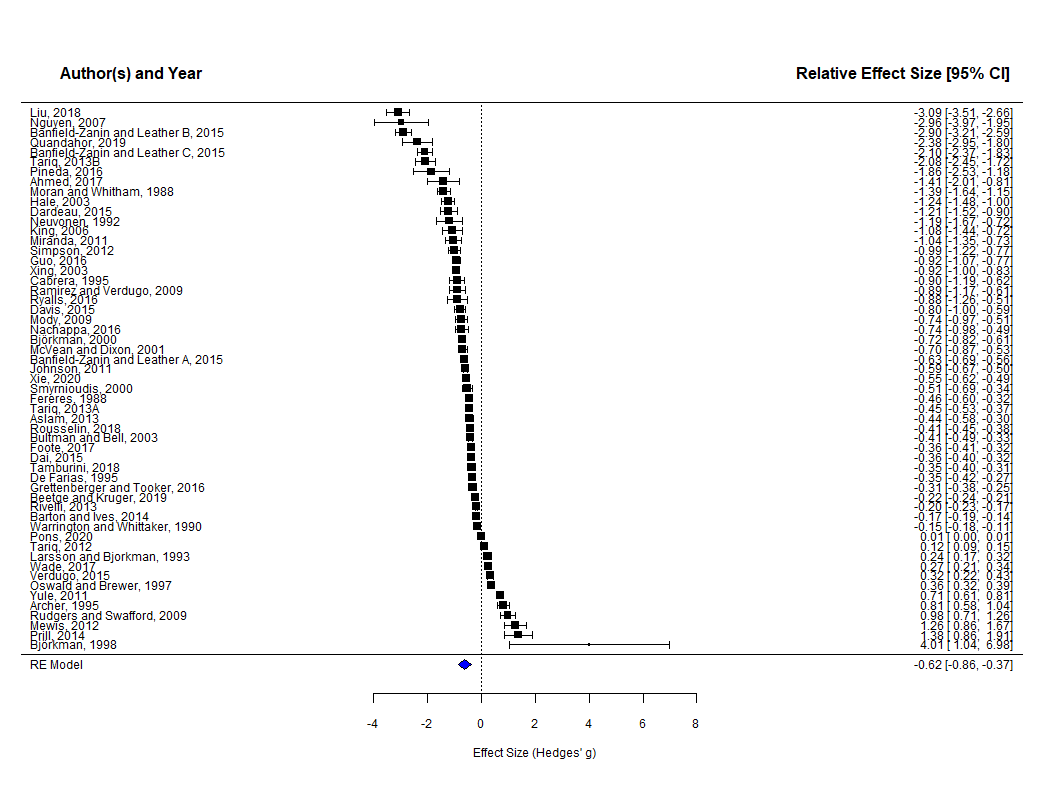
**Supplementary Fig. 4:** Forest plot of the 55 studies included in the meta-analysis of aphid fitness responses to plant drought stress. Plot displays the mean effect size and 95% confidence intervals of pooled aphid responses to drought stress. Blue diamond represents the relative effect size of the model.

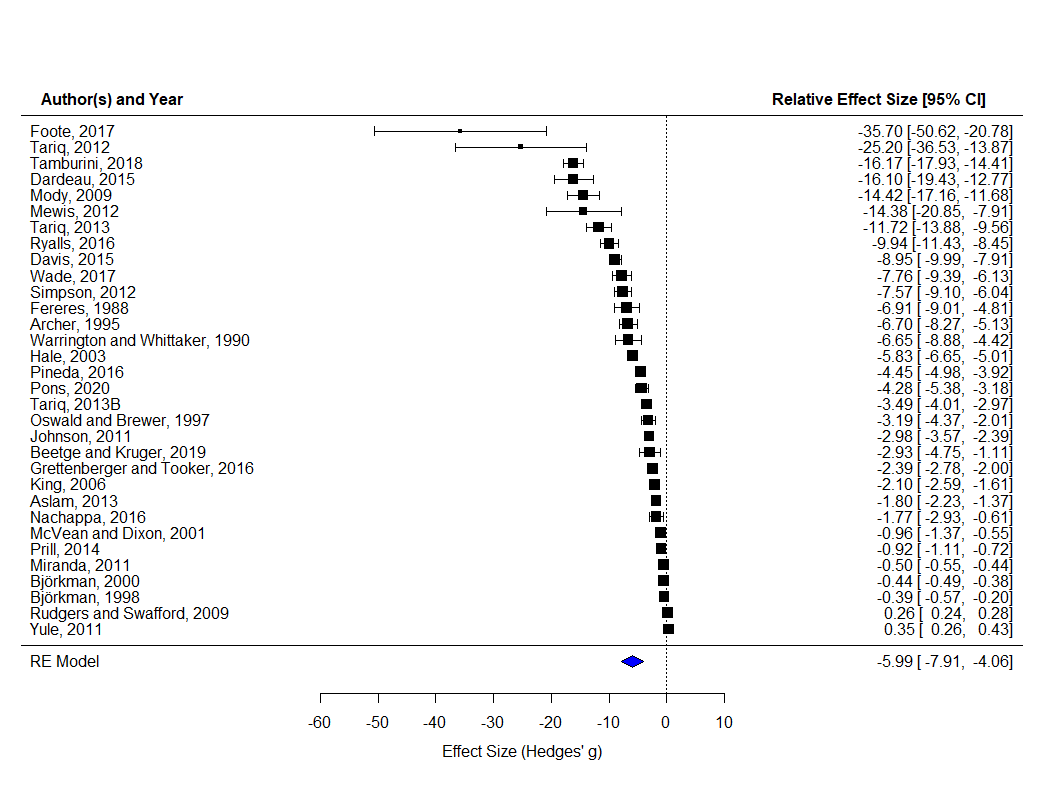
**Supplementary Fig 5:** Forest plot of the 32 studies included in the meta-analysis of plant physiological responses to drought stress. Plot displays the mean effect size and 95% confidence intervals. Blue diamond represents the relative effect size of the model.

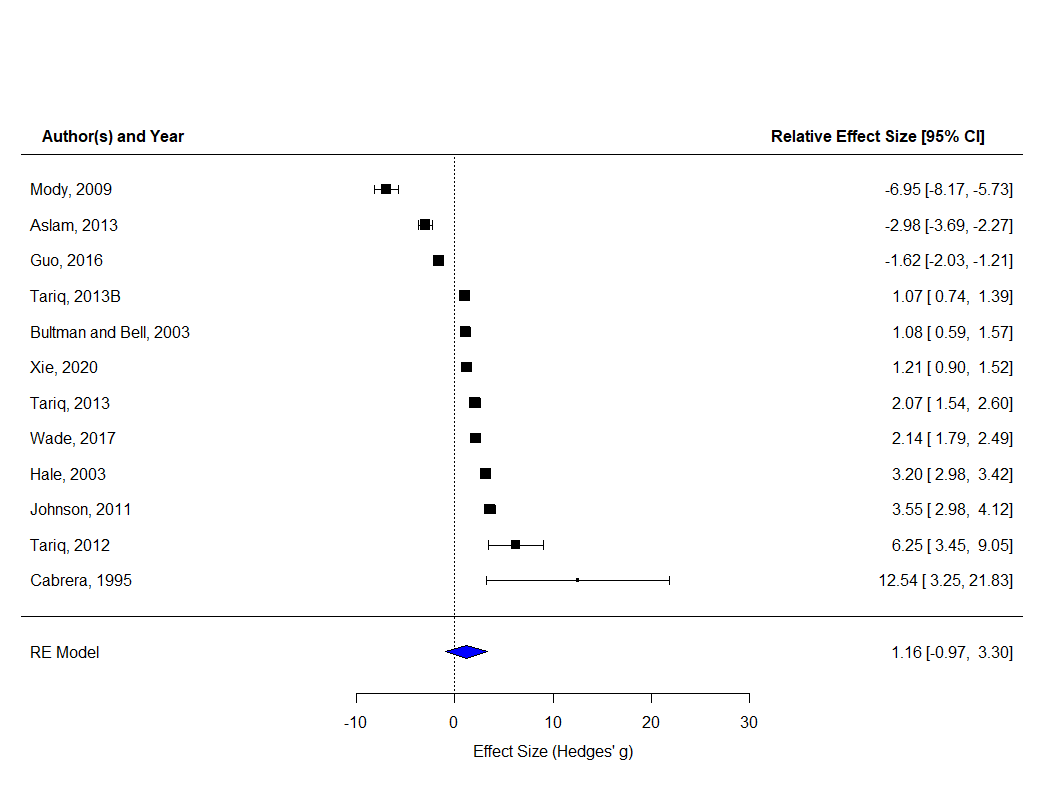

**Supplementary Fig 6:** Forest plot of the 12 studies included in the meta-analysis of plant nutritional responses to drought stress. Plot displays the mean effect size and 95% confidence intervals. Blue diamond represents the relative effect size of the model.

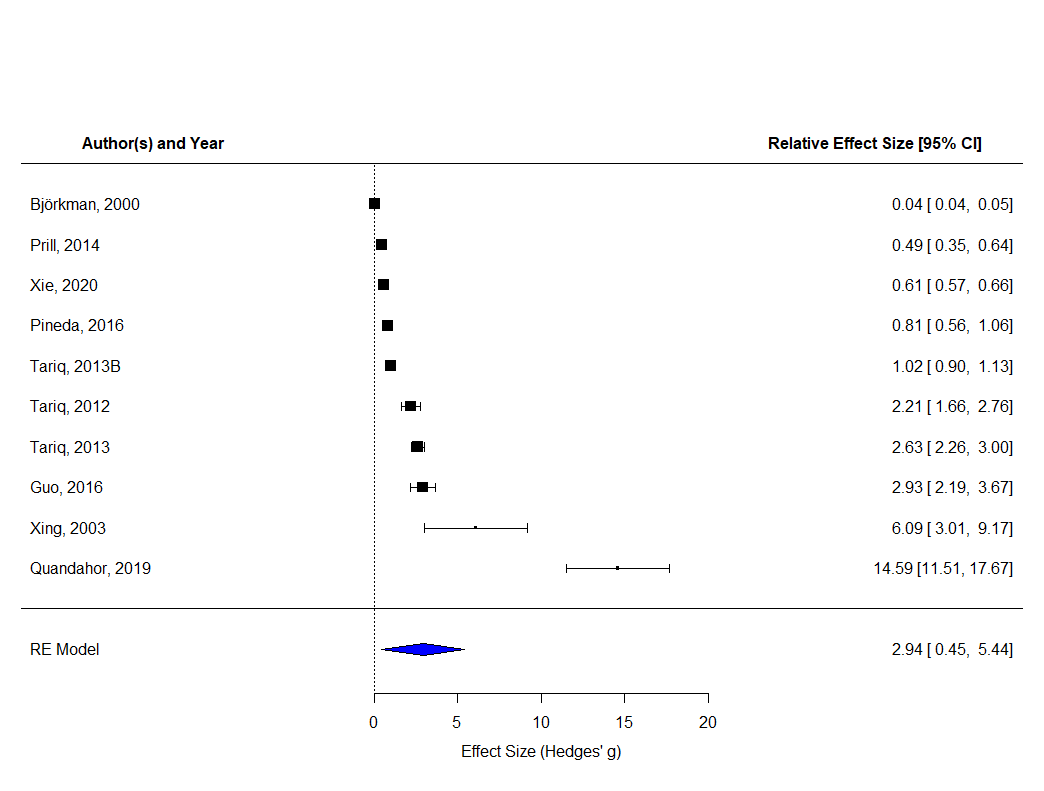
**Supplementary Fig 7:** Forest plot of the 7 studies included in the meta-analysis of plant defensive responses to drought stress. Plot displays the mean effect size and 95% confidence intervals. Blue diamond represents the relative effect size of the model.

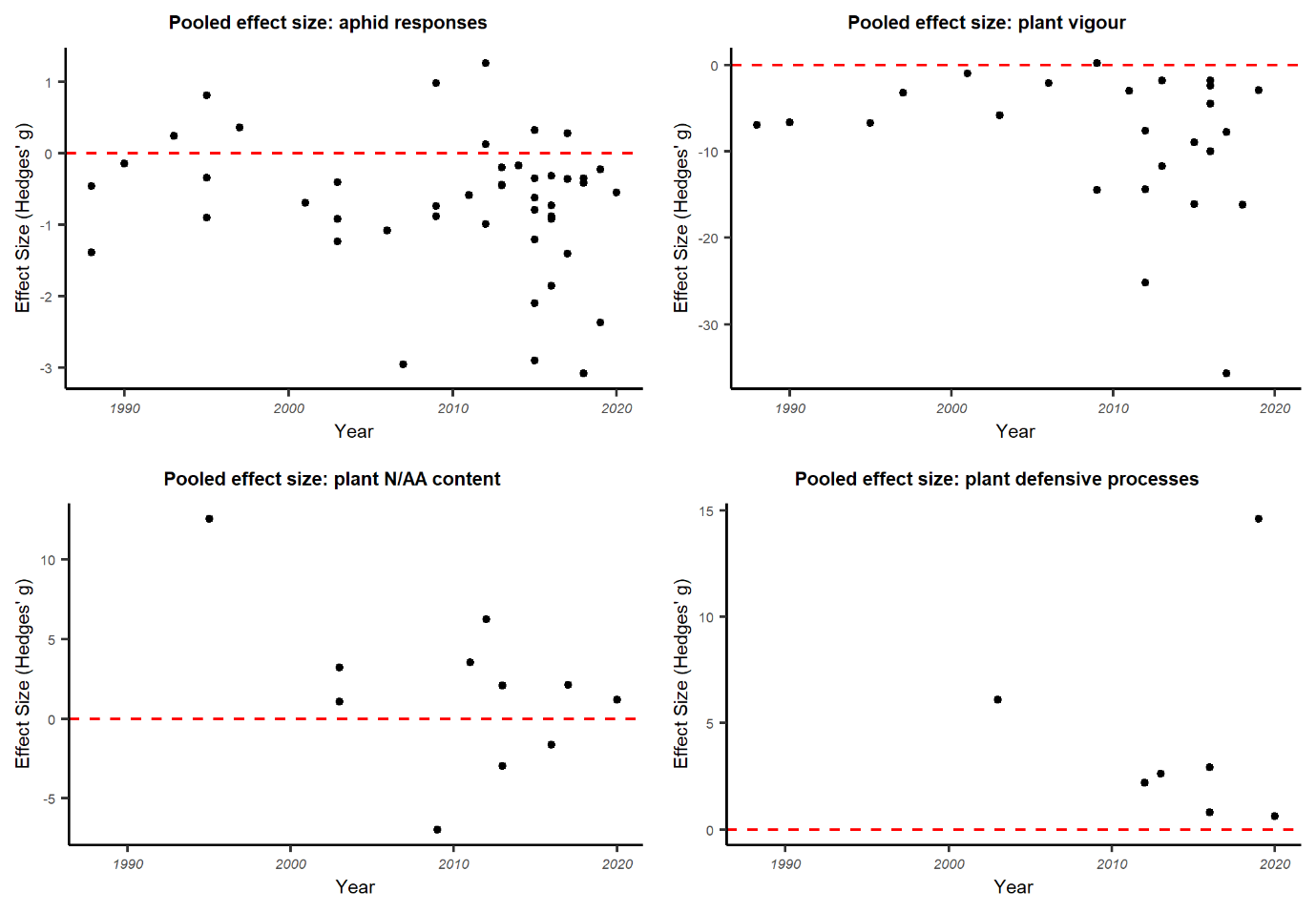
**Supplementary Fig 8:** Scatter plots showing aphid and plant responses to drought stress over publication time. Red dashed line represents zero effect size.

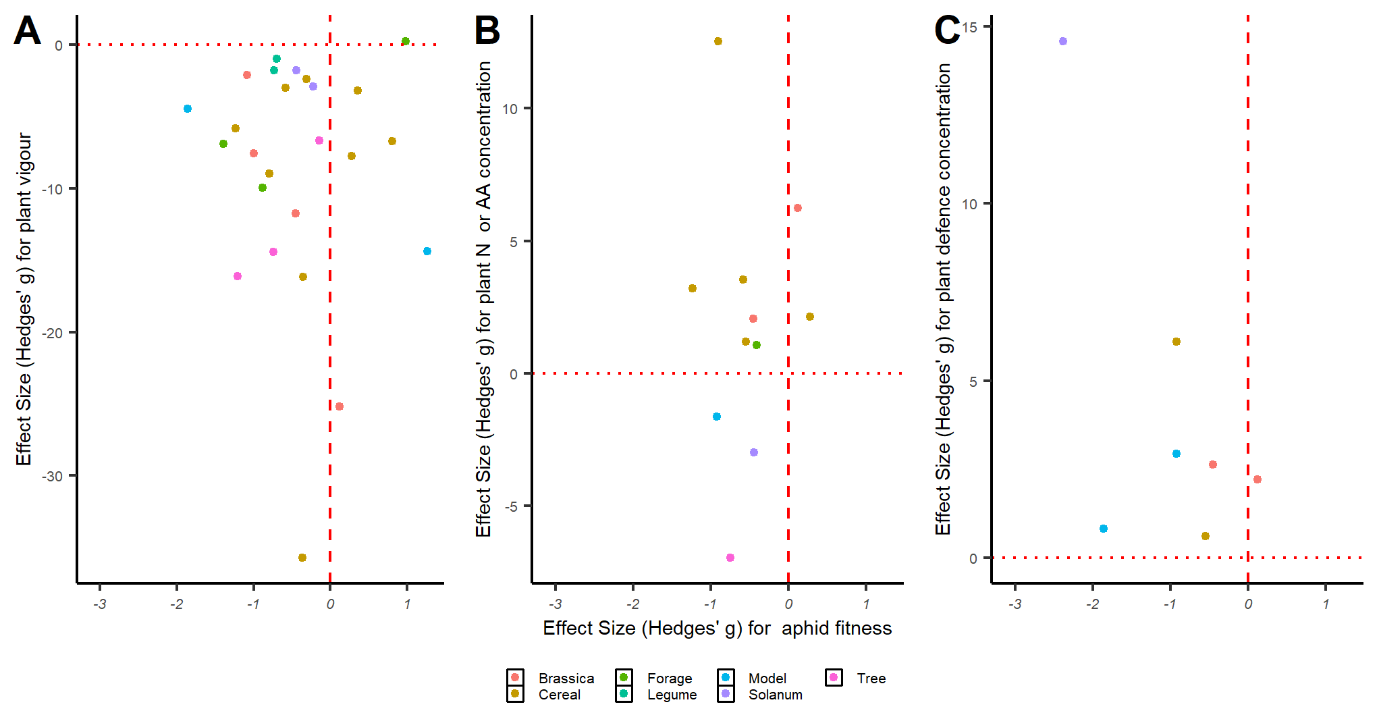

**Supplementary Fig 9:** Scatter plots showing the relationship between aphid and plant responses to drought stress. A: The relationship between plant vigour and aphid fitness, B: The relationship between plant N or AA concentration and aphid fitness, C: The relationship between plant defensive compound concentrations and aphid fitness. Red dashed line represents zero effect size for aphid fitness with the dotted red line representing the zero effect size for plant data (vigour, nutritional, defensive).
